# Repurposing Regulatory Toxicology Safety Data to Identify Potential Pro-Longevity Substances

**DOI:** 10.1101/2025.01.30.635304

**Authors:** Satomi Miwa, Olena Kucheryavenko, Kristin Herrmann, Christopher Saunter, Adelaide Raimundo, David Weinkove, Lars Niemann, Thomas von Zglinicki

**Author notes:** Corresponding authors Address for correspondence: Prof. Thomas von Zglinicki, Ageing Research Laboratories, Biosciences Institute, Faculty of Medical Sciences, Newcastle University, Newcastle upon Tyne NE4 5PL, UK., Satomi Miwa, Biosciences Institute, Faculty of Medical Sciences, Newcastle University, Newcastle upon Tyne NE4 5PL, UK. shared first authors (Satomi Miwa, Olena Kucheryavenko).

## Abstract

The repurposing of existing biosafety datasets offers unique opportunities in biomedical research. Here, we demonstrate how pesticide toxicity data, which include long-term survival studies in mammalian models, can be harnessed to uncover potential drug candidates or drug targets that can improve survival. We show that these substances frequently affect mitochondrial bioenergetics and that they can improve animal healthspan.

## Main text

Legally, pesticides must undergo rigorous *in vivo* safety testing before authorisation, including long-term toxicity and carcinogenicity assessments in two rodent species where survival is a primary endpoint^1^. These studies are subject to stringent quality controls, with findings stored by regulatory bodies such as the European Food Safety Authority (EFSA) in Europe and the Environmental Protection Agency (EPA) in the USA. We mined the electronic Archive of Studies and Assessments of the German Federal Institute for Risk Assessment (BfR), which contains toxicity data on approximately 1000 substances including long-term studies, testing multiple (mostly 3) doses in at least 50 rats and mice per dose and sex. Typically, long-term studies in rats lasted over two years, yielding median survival at endpoint in controls of 52% in males and 48% in females. In mice, only 33% of studies were conducted over 22 months, with similar median survival (57% in males and 52% in females); the remaining studies were shorter, resulting in higher survival and reducing statistical power to detect survival changes.

From 545 randomly taken database entries (Supp. Fig. S1), 154 contained long-term toxicity and survival data. Among these, 27 substances (18%) increased survival at study termination (P<0.05) in at least one species, sex, or dose. Only one, ascorbic acid, had previously been shown to extend lifespan in male mice^2^. Many substances also exhibited non-significant survival increases (10% or more) across various cohorts and concentrations, with survival decreases being rare (Table 1).

**Tab. 1:** 15 substances that increase rodent survival. All doses (given as ppm in food) that result in >10% change in survival at study termination for rats (R) or mice (M) are shown. Values in black indicate improved survival, red denotes decreased survival. *P<0.05, **P<0.01, ***P<0.005 and ****p<0.001 according to either Kaplan-Meier, Cox’s regression or trend test followed by pair-wise comparison. Changes in food intake (FI) and body weight (BW) are indicated by arrows if differences to controls were >10% over most of the test period.

| Substance | Dose range (ppm) | Males |  |  | Females |  |  |
| --- | --- | --- | --- | --- | --- | --- | --- |
|  |  | Mortality | FI | BW | Mortality | FI | BW |
| **Alanycarb (ALAN)** | R:30-300<br>M:100-1000 | R:30<br>M:100<br>M:300* |  |  | R:300<br>M:300<br>M:1000 |  |  |
| **Ascorbic acid (ASC)** | 25000,50000 | R:50000<br>M:25000<br>M:50000** |  |  |  |  |  |
| **Bifenazate (BIF)** | R:20-200<br>M:10-225 | R:80*<br>R:200*<br>M:100<br>M:225 | ↓ | ↓<br>↓<br>↓ |  |  |  |
| **Bromoxynil (BROMOXY)** | R:60-600<br>M:10-100 | R:60<br>R:600<br>M:30 |  | ↓ | R:60<br>R:190**<br>R:600***<br>M:10<br>M:30<br>M:100 |  | ↓<br>↓<br>↓ |
| **Bromuconazole (BROMUC)** | R:20-2000<br>M:20-500 | R:1000<br>R:2000<br>M:100***<br>M:500 |  |  | M:100<br>M:500 |  |  |
| **Chlorfenapyr (CHLORFE)** | R:60-600<br>M:20-240 |  |  |  | R:600*<br>M:240* | ↓<br>↓ | ↓<br>↓ |
| **Chlortoluron (CHLORTO)** | R:100-2500<br>M:100-2500 | R:100*<br>M:100*<br>M:500 |  |  | R:100*<br>R:500*<br>R:2500* | ↓ | ↓ |
| **Clothianidin (CLOTH)** | R:150-3000<br>M:100-2000 | R:1500****<br>R:3000****<br>M:100<br>M:1250 | ↓ | ↓<br>↓ | R:1500<br>R:3000***<br>M:350<br>M:1800* | ↓<br>↓<br>↓ | ↓<br>↓<br>↓ |
| **Cyantraniliprole (CYANT)** | R:20-20000<br>M:20-7000 | R:20<br>M:20*<br>M:150* |  |  | R:20<br>R:2000<br>M:7000 | ↓ | ↓ |
| **Cyproconazole (CYPROCON)** | R:20-350<br>M:5-200 | R:350<br>M:15*<br>M:100*<br>M:200**** |  | ↓<br>↓ | M:100****<br>M:200 |  | ↓<br>↓ |
| **Cyprodinil (CYPRODIN)** | R:5-2000<br>M:10-5000 | M:5000* |  | ↓ | R:75<br>M:5000 | ↓ |  |
| **Dimethenamid (DIMET)** | R:100-1500<br>M:30-3000 | R:700<br>R:1500*<br>M:30<br>M:300 | ↓ |  | R:1500<br>M:30<br>M:300<br>M:3000 |  | ↓ |
|  |  | M:3000 | ↓ |  |  |  |  |
| **Etoxazole (ETOX)** | R:50-10000<br>M:2250,4500 | R:10000<br>M:2250 | ? | ↓ | R:10000* | ↓ | ↓ |
| **Fenpyroximate (FEN)** | R:10-150<br>M:25-800 | R:25<br>R:75<br>R:150<br>M:400 | ↓<br>↓ | ↓<br>↓ | R:25<br>R:75*<br>R:150<br>M:100 | ↓<br>↓ | ↓<br>↓<br>↓ |
| **Lenacil (LEN)** | R:250-25000<br>M:100-7000 | R:2500*<br>R:25000<br>M:2500 |  | ↓ | R:250<br>R:25000<br>M:100<br>M:7000 |  | ↓ |

Out of these, 15 substances can be publicly disclosed for further investigation. 4 (alanycarb, ascorbic acid, bifenazate, bromuconazole) increased survival exclusively in males, while 2 (chlorfenapyr, chlortoluron) were only effective in females. Sex-biases for the remaining 9 substances were less pronounced (Tab. 1). Supp. Fig S2. provides examples of survival curves for bromoxynil.

Chlorfenapyr, chlothianidin and etoxazole tended to reduce food intake and body weight at the doses that improved survival (Tab. 1), suggesting dietary restriction might have played a role. In contrast, bromoxynil, cyproconazol and cyprodinil reduced body weight without reducing food intake albeit sex-specifically, implying they could have acted as ‘exercise mimetics’, or produced organ toxicity impacting on metabolism and/or the intestinal microbiome.

The most common chronic toxic effect, at least at maximum dose tested, was increased liver weight, often with centrilobular hypertrophy, indicating a detoxification response. None were regarded as genotoxic. Additional chronic toxicity indicators are detailed in suppl. Tab. S1. While adverse effects may not entirely disqualify these substances, a safety margin between beneficial and toxic doses is critical.

How might these substances promote mammalian survival? Perturbations of electron transfer systems were frequent mode of actions, although mostly in non-mammalian systems (suppl. Tab S2). Given that perturbations of mitochondria can activate stress response pathways and extend lifespan in *C. elegans* and mice ^3,4^, the effects of these 15 substances on cellular oxygen consumption rates (OCR) in mammalian cells were examined.

Multiple substances significantly inhibited intact cell OCR at high doses (LD80, Fig. 1; see suppl. Fig S3 for cell viability), but only bifenazate and fenpyroximate did so at LD20. Chlortoluron, cyprodinil, bromuconazole and bifenazate inhibited complex I exclusively, whereas etoxazole and fenpyroxymate also acted on additional sites in permeabilised cell experiments (Suppl. Fig. S4). On the other hand, bromoxynil and chlorfenapyr stimulated intact cell OCR, at both LD20 and LD80 for the former, and only at LD80 for the latter. Mitochondrial uncoupler activity of bromoxynil was seen in permeabilised cells in active respiration state (state 3, Suppl. Fig. S4) and was confirmed in resting state (state 4; suppl. Fig. S5) in line with the *in vivo* observation showing body weight reduction without changes in food intake (Table 1). Chlorfenapyr is a pro-drug to the active uncoupler, AC 303,268 (tralopyril) which typically requires activation by cytosolic oxidases^5^, and thus had no uncoupling activity in permeabilised cells (Suppl. Fig. S5). Therefore, bifenazate, bromoxynil, etoxazole and fenpyroxymate (and chlorfenapyr) may promote survival through perturbation of mitochondrial function (summarised in suppl. Tab S2) inducing pro-survival stress response pathways. These substances will, however, have other functions which could also be explored further as additional mechanisms of action.

**Fig. 1:**
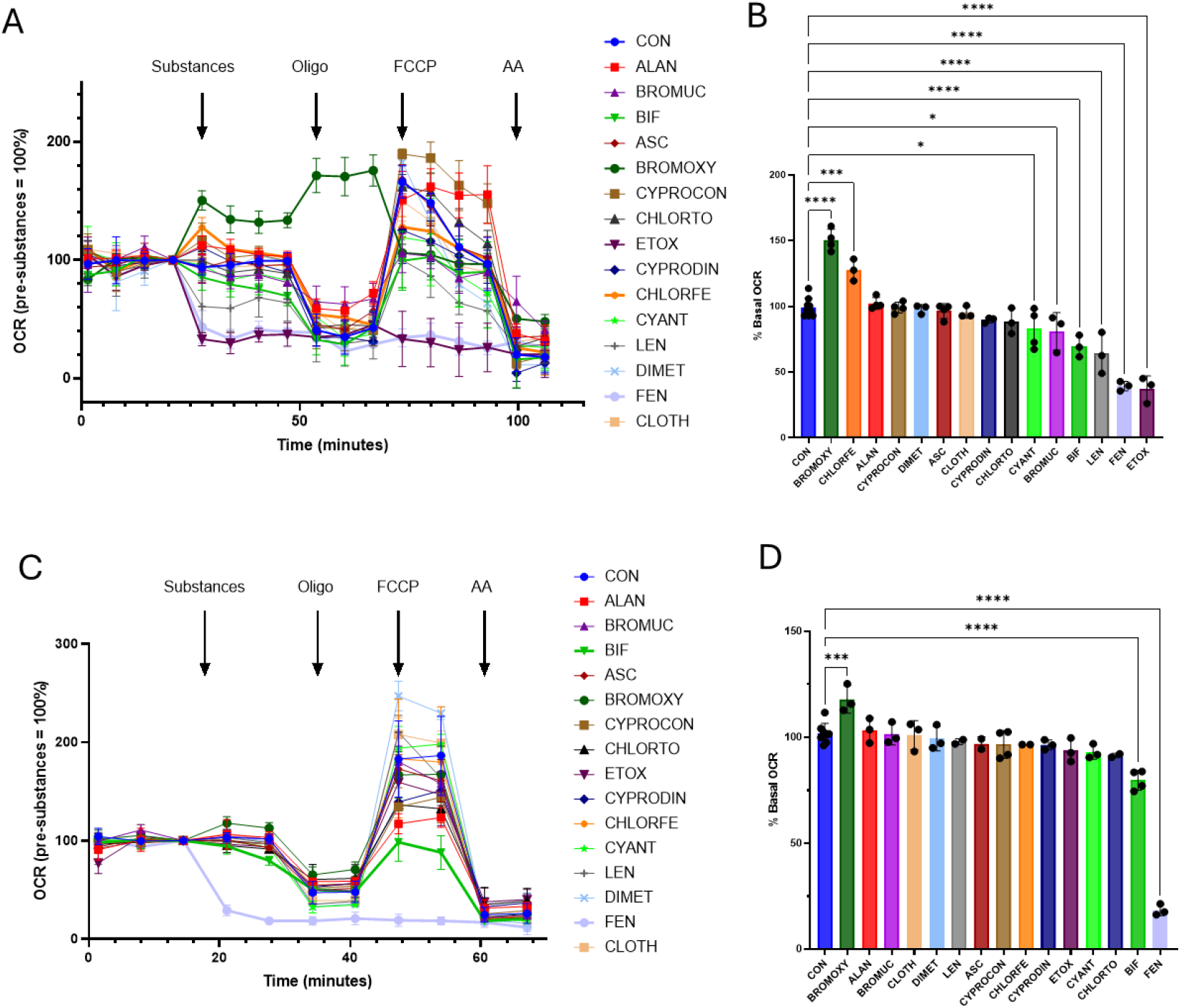
Effects of 15 substances at LD80 (A,B) and LD20 (C,D) on intact cell oxygen consumption rates (OCR) in human fibroblasts. A,C) Time course measurements of OCR. Sequential measurements of OCR at basal, substances at LD80 **(A)** or LD20 **(C)** concentrations, with oligomycin (Oligo, 0.5μM), with uncoupler FCCP (2.5μM), and with Antimycin A (AA, 2.5μM) using a Seahorse XF Analyzer. Data integrated from 3-9 (LD80) or 3-8 (LD20) measurements in 2-3 independent experiments. **B,D) % OCR in the presence of substances at LD80 (B) or LD20 (D) relative to that in the absence**. One way ANOVA, differences to controls (CON) indicated by asterisks with * P<0.01, ***P<0.001, ****P<0.0001.

Could these substances extend organismal healthspan? In a back-translation approach, *C. elegans* healthspan was examined using spontaneous motility, with two reliable indicators^6^: the proportion of moving worms and their average movement speed. Bifenazate and bromoxynil were selected as proof-of-concept examples, with the former inhibiting and the latter uncoupling mitochondria at LD20. Bifenazate increased the fraction of moving worms at higher ages (Fig. 2A) but not their movement speed (Fig. 2B), increasing the total time adult worms spent moving (Fig. 2C) but not distance (Fig. 2D). Moreover, it decreased movement speed in younger animals (Fig. 2B, suppl. Fig. S6A), suggesting a negative health effect at the tested concentrations. In contrast, under bromoxynil more worms remained active across the lifespan (Fig. 2E), moving faster at higher ages (Fig. 2F, suppl. Fig. S6B), and adult worms moved both longer (Fig. 2G) and further (Fig. 2H), indicating bromoxynil increased healthspan in *C. elegans*.

**Fig. 2:**
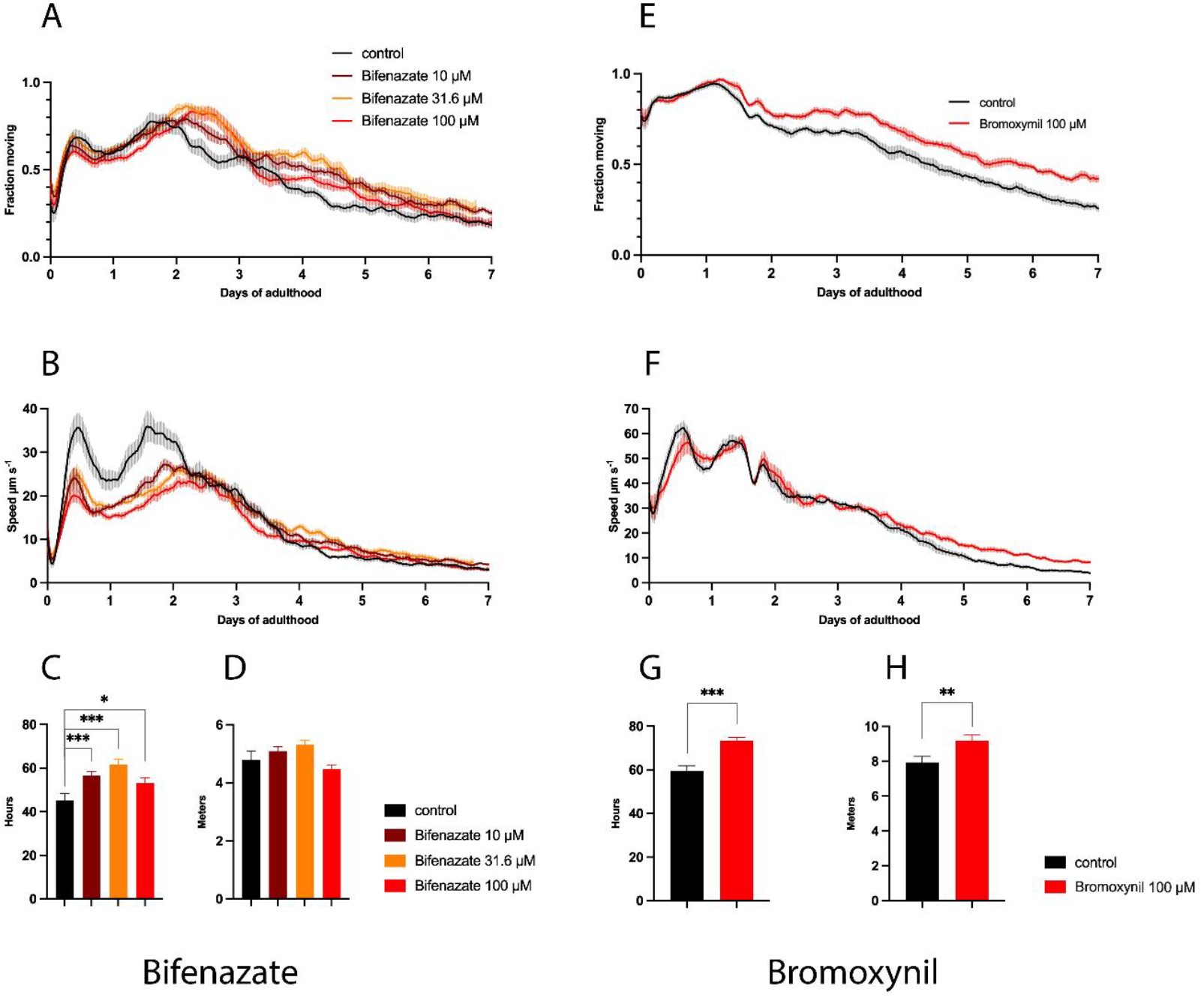
Effects of bifenazate (A-D) and bromoxynil (E-H) on healthspan indicators in *C. elegans*. **A,E)** Fraction of moving worms over their life history. **B,F)** Mean movement speed of all worms over time. Smoothed averages (bold lines) ± SEM (shaded areas), n≥350 animals per condition. **C,G)** Mean hours all worms moved between days 2 and 7. **D,H)** Mean distance travelled by worms between days 2 and 7. Means ± SEM over all worms from day 2 till 7, differences to controls indicated by asterisks with * P<0.05, **P<0.02, ***P<0.001.

In summary, our data mining efforts identified 15 out of 545 substances associated with greater survival in mammals for further study. Approximately half of them either inhibited or uncoupled mitochondrial electron transfer in mammalian cells, suggesting adaptive mechanisms linked with mitochondrial perturbations are key stress responses to support survival. Amongst them, bromoxynil, a mitochondrial uncoupler, was found to extend organismal healthspan in *C. elegans*, showing long-term toxicity studies, such as those submitted to national and international regulatory agencies, can be repurposed to identify new drugs or targets that could extend healthy lifespan.

## Acknowledgements

We thank Dr. Roland Solecki for support of this project and Dr. Michael Carroll for discussion.

## Funding

TvZ acknowledges funding from EU FP7-PEOPLE PITN-GA-2012-316964 (MARRIAGE) and H2020 WIDESPREAD Project 857524 (MIA Portugal). OK acknowledges funding from the Werner Baltes Fellowship, BfR.

## Data availability statement

Survival and toxicology data are in the Assessment Reports for the individual pesticides, available from the BfR or the EFSA upon request.

## Notes

### Competing Interest Statement

The authors have declared no competing interest.

