## Supplementary materials for "Repurposing Regulatory Toxicology Safety Data to Identify Potential Pro-Longevity Substances"

**Online Methods**

*Database screening*

The BfR electronic Archive of Studies and Assessments contains information about 1000 substances, covering a wide range of plant protection products (PPP) and co-formulants. Long-term survival data in males and females of two species (generally rats and mice) are available for about one third of the PPP. Unfortunately, automated electronic retrieval of survival data was not possible, therefore all survival data had to be manually retrieved. Quality of data was checked by a second independent observer. Screening strategy is shown in Fig. S1. Survival data for all concentrations of each drug per strain and sex were analysed by Kaplan-Meier product-limit method followed by Gehan-Breslow test and Holm-Sidak pairwise comparisons or Cox's regression model. Compounds for which survival data are in the public domain and that decreased mortality with $p<0.05$ in at least one species, one sex, and one dose are reported. As this was an exploratory analysis aimed at finding potential candidates for healthspan extension, we used the $P<0.05$ threshold without correction for multiple testing.

*Cell viability assay*

Human lung fibroblasts (MRC5) were seeded on a 96-well plate (15,000 cells per well), and the cells were treated with each of the 15 substances for 48 hours at concentrations ranging from 0.1µM to 5mM, except clothianidin, cyantraniliprol, dimethenamid, fenpyroximate and lenacil, where 50µM was the highest concentration, due to their poor solubility, and was used as 'pseudo LD80 concentration' if not reached at lower concentrations. Cell viability was then assessed using CellTiter-Glo® Luminescent Cell Viability Assay (Promega) according to manufacturer's instructions.

*Oxygen consumption rates (OCR) measurements*

Intact cell oxygen consumption rates (OCR) were measured using human lung fibroblasts (MRC5) in a Seahorse XF24 Analyzer (Agilent Technologies) in unbuffered Dulbecco's Modified Eagle Medium supplemented with 5mM glucose, 2mM L-glutamine and 3% FBS. The following compounds were added sequentially to test the effects of each of the 15 substances on cellular OCR: 0.5µM oligomycin to inhibit ATP synthase (hence inhibiting mitochondrial ATP generation), 2.5µM carbonyl cyanide p-trifluoromethoxy-phenylhydrazone (FCCP) to stimulate OCR, and 2.5µM Antimycin A to inhibit complex III.

For mechanistic studies, mitochondrial function was directly assessed by permeabilizing the cells with XF Plasma Membrane Permeabilizer (Agilent Technologies) according to the manufacturer's instructions, and mitochondria-specific OCR were determined in a Seahorse XF24 Analyzer. Initially, complex I-linked substrate, pyruvate (10mM) and malate (1mM) was supplemented and state 3 OCR was achieved by addition of 4mM ADP. Each of the 15 substances was then added to examine its complex I specific effect; state 3 OCR was chosen to sensitively test any inhibitory effects of the substances. Rotenone (5µM) was added next to inhibit complex I, followed by the complex II-linked substrate succinate (4mM) to test the effects of each of the substances on other sites than complex I. To specifically test mitochondrial uncoupling function of bromoxynil and chlorfenapyr, permeabilised cell experiments were conducted as above, but their effects were tested at state 4 condition (in the presence of oligomycin; using complex I-linked substrate), along known uncouplers, FCCP, niclosamide and BAM15 as positive controls.

*C. elegans healthspan determination*

Worms to be tested (SS104 *glp-4(bn2)* obtained from the *Caenorhabitis* Genetics Center, University of Minnesota) were maintained at 15°C. For healthspan testing, 3 days after a timed egg lay worms were shifted to 24°C. The following day worms at the L4 stage were transferred onto agar Petri dishes containing the substances to be tested and controls. Movement of the population of worms was monitored from the L4 stage onwards using the WormGazer$^{TM}$ technology as described [1]. Outputs were the fraction of the population moving above a threshold speed and the mean speed of those worms. The mean speed of all worms was obtained by multiplying those parameters together. Statistical analysis was done as described [1].

**Supplementary Tables**

**Tab. S1. Toxicity summary for 15 substances that improve rodent survival in long-term studies.** Data obtained from the BfR Archive of Studies and Assessments. Numbers indicate either substance concentrations in food (ppm) or doses directly applied by oral gavage (in mg/kg bw per day). NOAEL: No Observed Adverse Effect Level, LOAEL: Lowest Dose with Observed Adverse Effect, MoA: Mode of action. Current classification obtained from either the website of European Chemicals Agency (ECHA, https://www.echa.europa.eu/information-on-chemicals, retrieved in December, 2024) or from proposals of the European Food Safety Authority (EFSA).

| Substance / Toxicological endpoint | Genotoxicity | Reproductive toxicity (incl. developmental effects) | Neurotoxicity | Carcinogenicity | Chronic (organ) toxicity | Toxicity classification (oral route*) |
|---|---|---|---|---|---|---|
| **Alanycarb** | Negative | Reduced pup weight from 120 ppm onwards and lower pup survival at 480 ppm in a multi-generation study, NOAEL 30 ppm (rats) | Signs of cholinesterase inhibition at toxic doses | No indication | Nervous system (cholinesterases↓), liver (organ weight↑), blood (some evidence of anaemia); NOAELs 30 ppm in rats, 100 ppm in mice | Not listed by ECHA, no proposals by EFSA |
| **Ascorbic acid** | Negative | No indication | No indication | No indication | No indications up to highest tested doses of ca 2000 mg/kg bw per day (based on old studies in rats and mice) | None |
| **Bifenazate** | Negative | Lower body weight and some delay in female sexual maturation in a two-generation study | Weak evidence; NOAEL 13 mg/kg bw per day, based on reduced | No indication | Spleen, kidney and liver findings (organ weights, histopathology), anaemia; NOAELs 20 ppm in rats, 10 ppm in mice | STOT RE 2, H373 |

| | | | | | | |
|---|---|---|---|---|---|---|
| | | on rats, NOAEL 80 ppm; variations in development of vascular system in rat foetuses, NOAEL 100 mg/kg bw per day | rearing at 35 mg/kg bw per day (rat) | | | |
| **Bromoxynil** | Negative | Reduced bodyweight gain in offspring in a multi-generation study in rats, NOAEL 50 ppm; fetal malformations at a high (maternally toxic) dose of 40 mg/kg bw per day (rats) | No indication | Hepatocellular adenoma and carcinoma in male mice at 20-30 ppm and above and at 300 ppm in females but not relevant for humans due to MoA | Liver (organ weight↑, centrilobular hypertrophy, spongiosis hepatis), thyroid (reduced hormone levels, histopathology); NOAELs 60 ppm in rats, 30 ppm in mice | Repr. 2, H61d; Acute Tox. 3, H301 |
| **Bromuconazole** | Negative | Lower body weight gain in offspring, NOAEL 20 ppm (rat); increase in placental weight, ossification delay and skeletal variations in rat foetuses, NOAEL 10 mg/kg bw per day | No indication | Hepatocellular carcinoma at top dose levels of 2000 or 3000 ppm in rats and mice, probably not relevant to humans | Liver (organ weight↑, histopathology, clinical chemistry), adrenals (vacuolation); NOAELs 20 ppm (both rat and mouse) | Not listed by ECHA; EFSA proposal: Acute Tox. 4, H302; Repr. 2, H361d |

| | | | | | | |
|---|---|---|---|---|---|---|
| **Chlorfenapyr** | Negative | Reduced pup survival in rats during postnatal days 1-4; tentative NOAEL 60 ppm | Vacuolation of white matter in the brain, spinal cord and optic nerve; NOAEL 60 ppm (rat) | No indication | Liver weight↑, red blood cell parameters↓; NOAELs 60 ppm in rats, 20 ppm in mice | Acute Tox. 4, H302 |
| **Chlorotoluron** | Negative | Resorption rate↑ and some evidence of skeletal anomalies in rabbits, NOAEL 50 mg/kg bw per day; lower number of implantation sites in dams in a two-generation study in rats, NOAEL 1000 ppm | No indication | Kidney tumours↑ and positive trend for liver tumours in male mice at 2500 ppm, NOAEL 500 ppm | Kidney weight↓, some evidence of anaemia; NOAEL 100 ppm (both in rats and mice) | Carc. 2, H351; Repr. 2, H361d |
| **Clothianidin** | Mixed results in vitro but overall negative (based on in vivo studies) | Low sperm motility in parental males, delayed sexual maturation in offspring in a two-generation study in rats, NOAEL 150 ppm; visceral anomalies and reduced | Limited evidence of reduced activity and neuro-developmental effects, perhaps due to systemic toxicity | No indication | Kidney (hyperplasia, mineralisation), liver (eosinophilic foci and hypertrophy), ovaries (interstitial cell hyperplasia), myocardial degeneration; NOAELs 150 ppm (rat), 350 ppm (mouse) | Acute Tox. 4, H302 |

| | | | | | | |
|---|---|---|---|---|---|---|
| | | ossification in rabbit foetuses, NOAEL 25 mg/kg bw per day | | | | |
| **Cyantraniliprole** | Negative | No indication | No indication | No indication | Liver (organ weight↑, histopathology, clinical chemistry); NOAELs 200 pm in rats, 150 ppm in mice | None |
| **Cyproconazole** | Negative | Malformations in rat and rabbit foetuses; NOAELs 2 mg/kg bw per day (rabbit), 12 mg/kg bw per day (rat) | No indication | Liver cell adenomas and carcinomas in male mice, human relevance equivocal; NOAEL 15 ppm | Liver (histopathology, clinical chemistry, organ weight↑); NOAELs 50 ppm in rats, 15 ppm in mice | Repr. 1B, H360D; STOT RE2, H373; Acute Tox. 3, H301; additional proposal by EFSA: Carc. 2, H351 |
| **Cyprodinil** | Negative | No indication | No indication | No indication | Liver (organ weight increase, histopathology), pancreas (hyperplasia of acinar cells); NOAEL 75 ppm in rats, 150 ppm in mice | None |
| **Dimethenamid** | Negative | Delayed ossification in rat foetuses, NOAEL 25 mg/kg bw per day | No indication | Equivocal increase in ovarian hyperplasia/adenoma in rats, NOAEL 700 ppm | Liver (organ weight↑ and histopathology); NOAELs 100 ppm (rat), 300 ppm (mouse) | Acute Tox. 4, H302 |
| **Etoxazole** | Negative | Increase in peri-/postnatal mortality of pups in a two-generation study | No indication | No indication | Liver (organ weight↑ and histopathology), effects in teeth and bone suggestive of fluorosis; NOAELs 50 | None |

| | | | | | | |
|---|---|---|---|---|---|---|
| | | in rats; NOAEL 400 ppm | | | ppm (rat), 2250 ppm (mouse) | |
| **Fenpyroximate** | Negative | No indication | No indication | No indication | Ovarian atrophy in mice; NOAELs (based on reductions in body weight and food consumption) 25 ppm in rats, 100 ppm in mice | Acute Tox. 3, H301 |
| **Lenacil** | Negative | Skeletal variations in rat foetuses↑ suggestive of developmental delay; NOAEL 100 mg/kg bw per day | No indication | Increase in mammary gland and uterine adenocarcinoma in female rats, NOAEL 250 ppm; liver adenoma and lung carcinoma in male mice, NOAEL:2500 ppm | Kidney, liver, spleen, and thyroid (organ weights↑, histopathology, clinical chemistry); NOAELs 250 ppm (rat) or 100 ppm (mouse) | Carc. 2, H351 |

*Any classification for endpoints such as acute dermal or inhalative toxicity, skin and eye irritation or skin sensitization was not taken into consideration for this table.

| Substance | Class | Mode of action | Reference | Our data | | | |
|---|---|---|---|---|---|---|---|
| | | | | Intact cell OCR (LD20) | Intact cell OCR (LD80) | Mitochondrial OCR: complex I-linked, LD80 | Mitochondrial OCR: complex II-linked, LD80 |
| Alanycarb | Insecticide | Cholinesterase activity inhibitor | 2 | | | | |
| Ascorbic acid | Fungicide, bacteriocide | Phytoalexin synthesis activator, water-soluble anti- or pro-oxidant | 3 | | | | |
| Bifenazate | Insecticide | GABA receptor agonist; ETC inhibitor | 4 | ↓ | ↓ | ↓ | |
| Bromoxynil | Herbicide | Photosynthetic ETC inhibitor; OXPHOS uncoupler | 5 | ↑ | ↑ | ↑ | ↑ |
| Bromuconazole | Fungicide | Ergosterol biosynthesis inhibitor; Fungal CYP inhibitor | 6 | | ↓ | ↓ | |
| Chlorfenapyr | Pro-insecticide | Pro-drug to the active uncoupler, AC 303,268 (tralopyril) | 7 | | ↑ | | |
| Chlortoluron | Fungicide | Photosynthetic ETC inhibitor | 8,9 | | | ↓ | |
| Clothianidin | Insecticide | Targets AChR, suppresses NF-kB signalling in the honey bee | 10 | | | | |
| Cyantraniliprole | Insecticide | Disrupts the $Ca^{2+}$ balance; Second generation ryanodine receptor | 11 | | ↓ | | |
| Cyproconazole | Fungicide | Cytochrome P450 inhibitor (CYP51) | 12 | | | | |
| Cyprodinil | Fungicide | Methionine biosynthesis inhibitor; COX2 - PDG2 inhibitor | 13 | | | ↓ | |
| Dimethenamid | Herbicide | Selective fatty acid synthesis inhibitor | 14 | | | | |
| Etoxazole | Insecticide | Inhibits chitin synthase 1, may enhance reactive oxygen species production | 15 16 | | ↓ | ↓ | ↓ |
| Fenpyroximate | Acaricide | Blocks complex I by binding to ND5 | 17 18 | ↓ | ↓ | ↓ | ↓ |

| | | | | | |
|---|---|---|---|---|---|
| Lenacil | Herbicide | Inhibits photosynthesis (photosystem II) | [19] | | ↓ |

**Tab. S2. Possible mechanisms of action of 15 substances as per literature and summary of their mitochondrial effects in human fibroblasts.** Red colour indicates that according to the literature, effects on electron transfer systems are indicated, albeit not necessarily in mammalian systems. Arrows indicate a significant effect of the substance in human fibroblasts. In mitochondrial OCR, where '↓' is shown, 'complex I-linked' indicates the inhibitory effects at complex I, whereas that for 'complex II-linked' are those other than at complex I. OCR, Oxygen Consumption Rates; ETC, electron transport chain; OXPHOS, oxidative phosphorylation.

**Supplementary Figures**

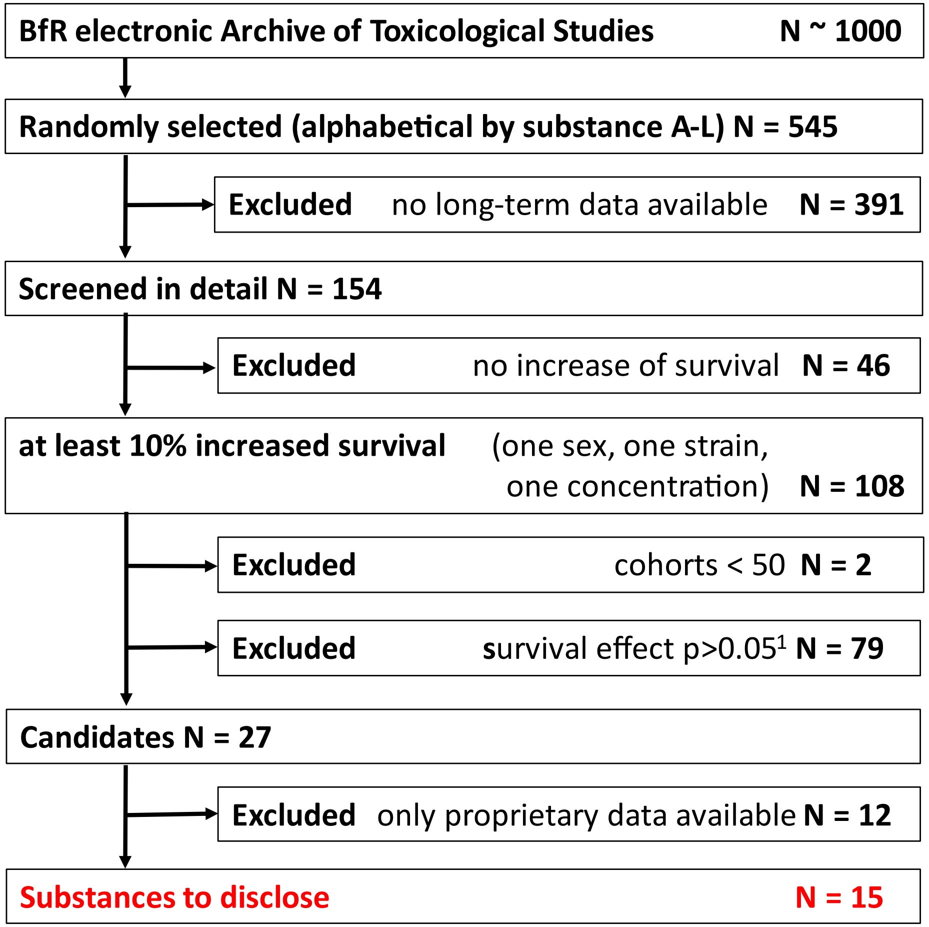

**Fig. S1: Screening strategy.** [1]Significance established by Cox test, Tarone test, or LogRank test.

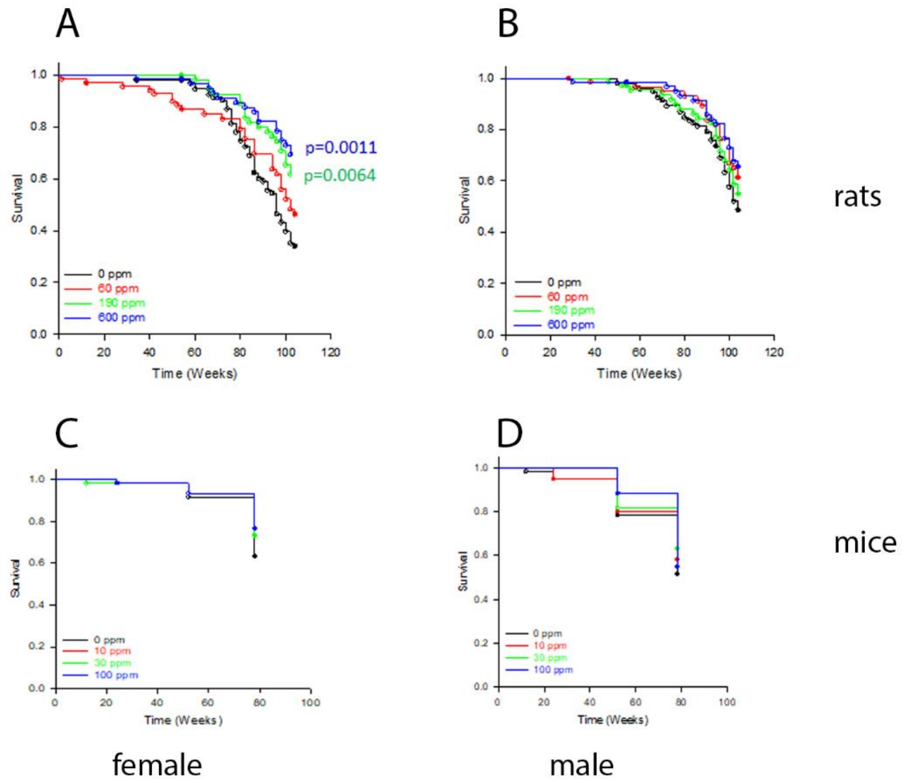

**Fig. S2: Example mortality curves for rats and mice treated with bromoxynil.** A) female rats, B) male rats, C) female mice, D) male mice. P values are given when significant according to Gehan-Breslow test and Holm-Sidak pairwise comparisons.

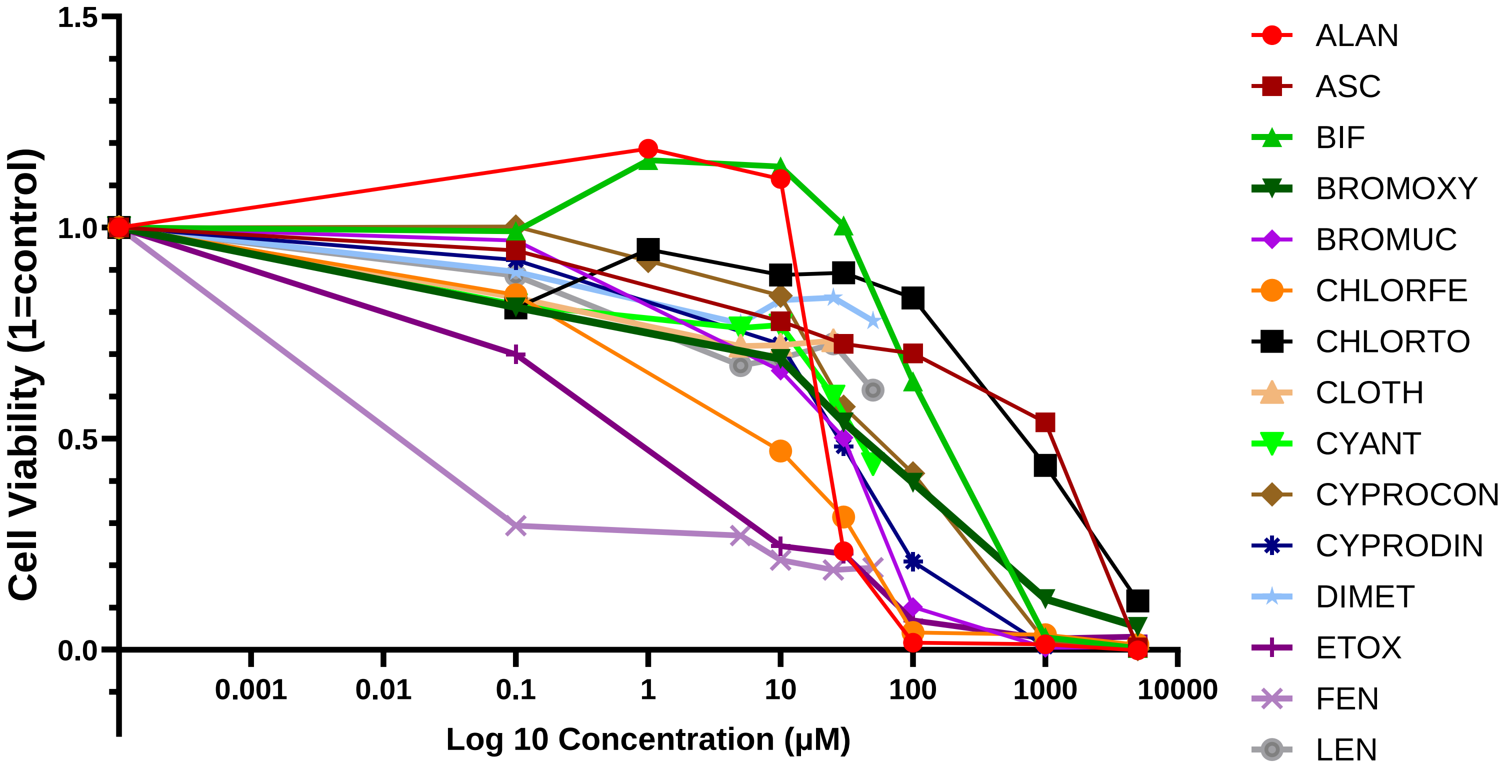

**Fig. S3: Human lung fibroblasts (MRC5) viability following 48h treatment with 15 selected substances at the indicated concentrations.** The maximum concentrations tested for clothianidin, cyantraniliprol, dimethenamid, fenpyroximate and lenacil were 50µM due to their poor solubility.

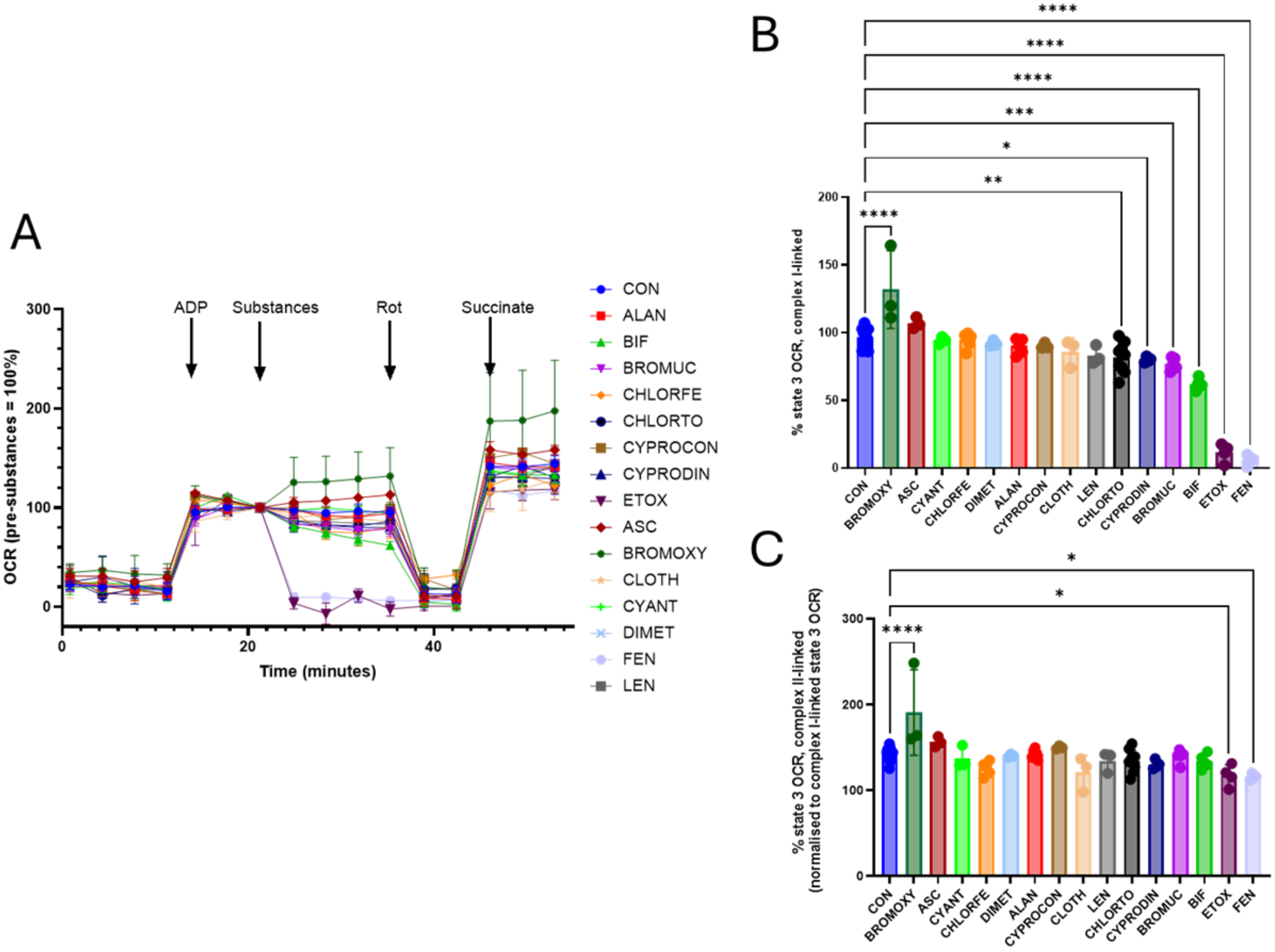
A
B
C
ADP
Substances
Rot
Succinate
OCR (pre-substances = 100%)
Time (minutes)
0
20
40
100
200
300
CON
ALAN
BIF
BROMUC
CHLORFE
CHLORTO
CYPROCON
CYPRODIN
ETOX
ASC
BROMOXY
CLOTH
CYANT
DIMET
FEN
LEN
% state 3 OCR, complex I-linked
0
50
100
150
200
****
**
*
***
****
****
****
CON
BROMOXY
ASC
CYANT
CHLORFE
DIMET
ALAN
CYPROCON
CLOTH
LEN
CHLORTO
CYPRODIN
BROMUC
BIF
ETOX
FEN
% state 3 OCR, complex II-linked
(normalised to complex I-linked state 3 OCR)
0
100
200
300
****
*
*
CON
BROMOXY
ASC
CYANT
CHLORFE
DIMET
ALAN
CYPROCON
CLOTH
LEN
CHLORTO
CYPRODIN
BROMUC
BIF
ETOX
FEN

**Fig. S4: Effects of 15 substances on complex I- and II- linked mitochondrial oxygen consumption rates (OCR) in permeabilised human lung fibroblasts. A)** Sequential measurements of mitochondrial OCR in the presence of complex I-linked substrates (Pyruvate 10mM and Malate 1mM) at basal, at State 3 (ADP 4mM), with substances at LD80 (see suppl. Fig. S3), with complex I inhibitor rotenone (5µM), and with the complex II substrate, succinate (4mM) added at the times indicated. **B)** % mitochondrial OCR at state 3 in the presence of complex I-linked substrate, with substances (at LD80) relative to that in the absence. **C)** % mitochondrial OCR at state 3 in the presence of complex II-linked substrate succinate, with substances (at LD80), relative to that in the absence (normalised to complex I-linked OCR, i.e. the OCR after ADP addition). Data from 3-14 measurements in 2-3 independent experiments using Seahorse XF Analyzer. One way ANOVA, differences to CON indicated by asterisks with * $P<0.01$, ***$P<0.001$, ****$P<0.0001$.

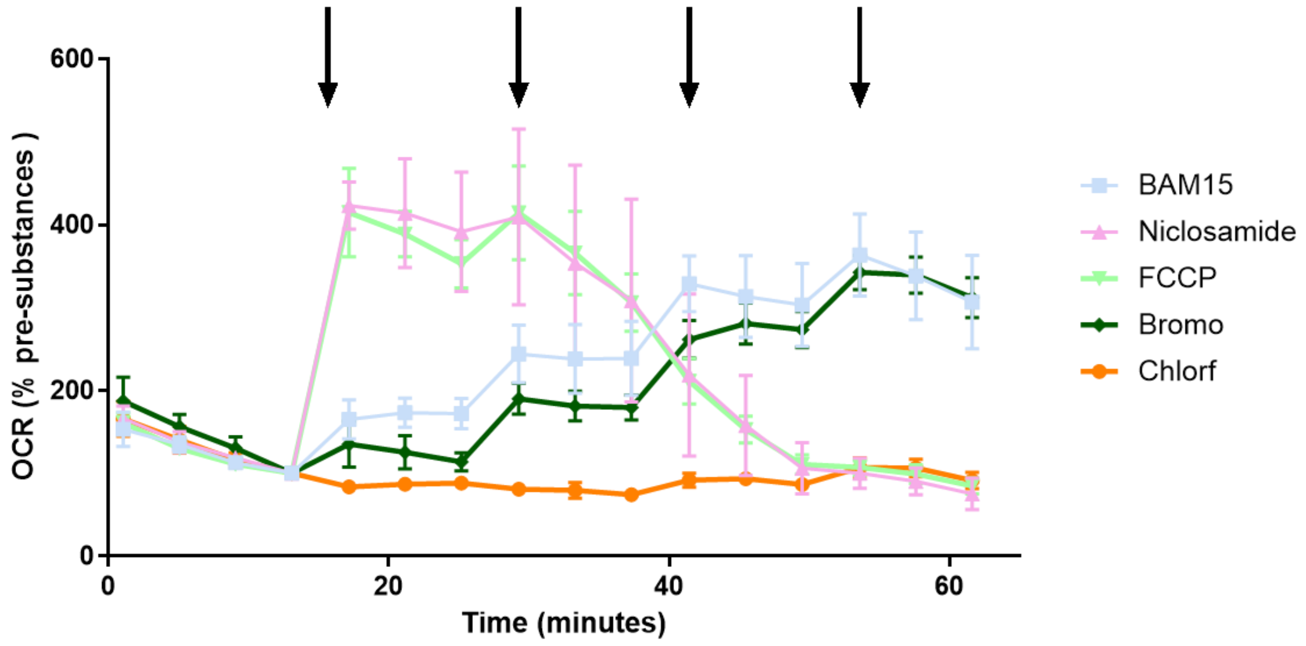

**Fig. S5: Mitochondrial OCR (at state 4) in permeabilised MRC5 cells, supplemented with complex I-linked substrates 10mM Pyruvate and 1mM Malate in the presence of 5µM oligomycin.** At each arrow, mitochondrial uncouplers BAM15, FCCP or Niclosamide at 1µM (final concentration after 4 additions = 4µM), or bromoxinil or chlorfenapyr at 10µM (final concentration after 4 additions = 40µM) were added.

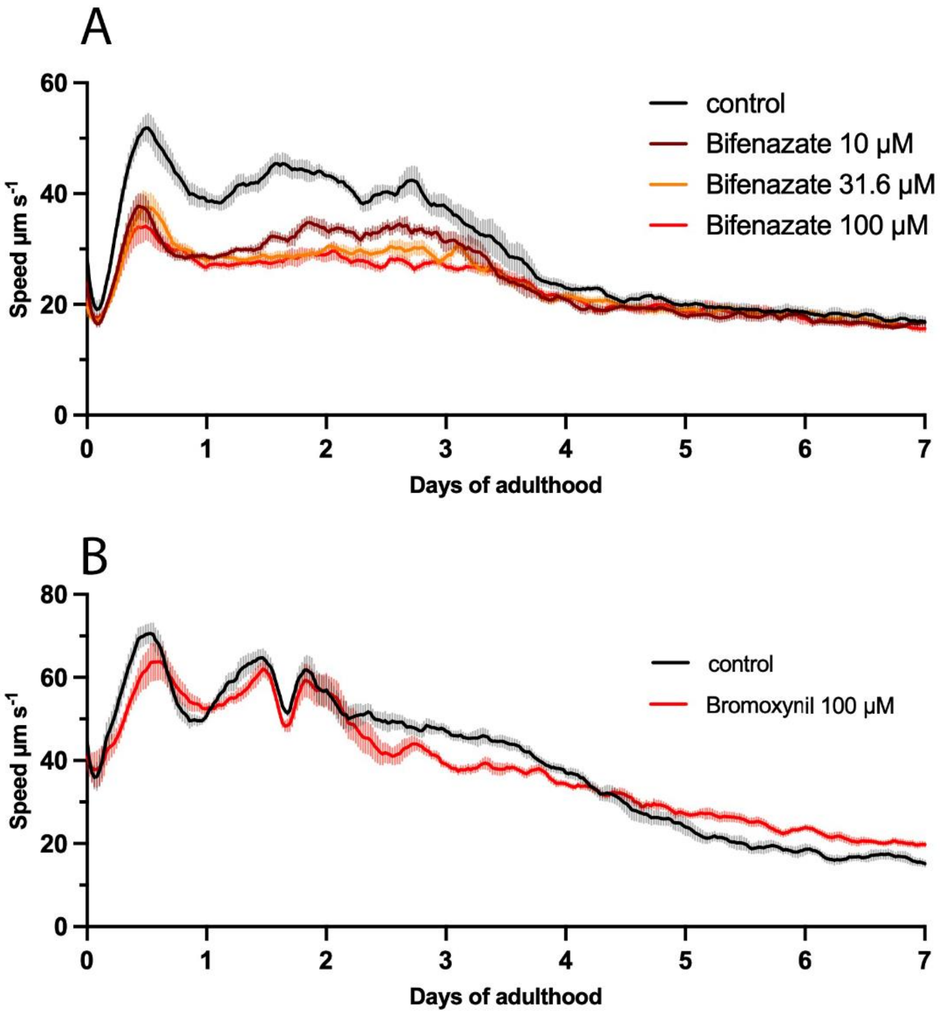

**Fig. S6: Mean movement speeds of moving worms (excluding non-moving animals) over their life history under bifenazate (A) or bromoxynil (B).** Data are smoothed averages (bold lines) ± SEM (shaded areas), n ≥ 350 animals per condition.

6 *Pesticide Properties Database: Bromuconazole (Ref: LS 860263)*, <https://sitem.herts.ac.uk/aeru/ppdb/en/Reports/97.htm> (

7 *US EPA Pesticide Fact Sheet: Chlorfenapyr* <https://www3.epa.gov/pesticides/chem_search/reg_actions/registration/fs_PC-129093_01-Jan-01.pdf> (

8 Broser, M. *et al.* Structural basis of cyanobacterial photosystem II Inhibition by the herbicide terbutryn. *J Biol Chem* **286**, 15964-15972 (2011). https://doi.org:10.1074/jbc.M110.215970

9 Diaz-Munoz, G., Miranda, I. L., Sartori, S. K., de Rezebde, D. C. & Diaz, M. A. N. in *Studies in Natural products Chemistry* Vol. 58 (ed Atta-ur-Rahman) 313-338 (Elsevier, 2018).

10 Di Prisco, G. *et al.* Neonicotinoid clothianidin adversely affects insect immunity and promotes replication of a viral pathogen in honey bees. *Proc Natl Acad Sci U S A* **110**, 18466-18471 (2013). https://doi.org:10.1073/pnas.1314923110

11 Sparks, T. C. *et al.* Insecticides, biologics and nematicides: Updates to IRAC's mode of action classification - a tool for resistance management. *Pestic Biochem Physiol* **167**, 104587 (2020). https://doi.org:10.1016/j.pestbp.2020.104587

12 He, Z. *et al.* Stereostructure-activity mechanism of cyproconazole by cytochrome P450 in rat liver microsomes: A combined experimental and computational study. *J Hazard Mater* **416**, 125764 (2021). https://doi.org:10.1016/j.jhazmat.2021.125764

13 Kugathas, S. *et al.* Effects of Common Pesticides on Prostaglandin D2 (PGD2) Inhibition in SC5 Mouse Sertoli Cells, Evidence of Binding at the COX-2 Active Site, and Implications for Endocrine Disruption. *Environ Health Perspect* **124**, 452-459 (2016). https://doi.org:10.1289/ehp.1409544

14 Lamberth, C. in *Bioactive Carboxylic Compound Classes* 293-302 (2016).

15 Demaeght, P. *et al.* High resolution genetic mapping uncovers chitin synthase-1 as the target-site of the structurally diverse mite growth inhibitors clofentezine, hexythiazox and etoxazole in Tetranychus urticae. *Insect Biochem Mol Biol* **51**, 52-61 (2014). https://doi.org:10.1016/j.ibmb.2014.05.004

16 Sun, D., Pang, J., Fang, Q., Zhou, Z. & Jiao, B. Stereoselective toxicity of etoxazole to MCF-7 cells and its dissipation behavior in citrus and soil. *Environ Sci Pollut Res Int* **23**, 24731-24738 (2016). https://doi.org:10.1007/s11356-016-7393-7

17 Boerner, H., Schlueter, K. & Aumann, J. *Pflanzenkrankheiten und Pflanzenschutz*. 8 edn,  (Springer, 2009).
18 Shiraishi, Y., Murai, M., Sakiyama, N., Ifuku, K. & Miyoshi, H. Fenpyroximate binds to the interface between PSST and 49 kDa subunits in mitochondrial NADH-ubiquinone oxidoreductase. *Biochemistry* **51**, 1953-1963 (2012). https://doi.org:10.1021/bi300047h
19 *PesticideProperties Database: Lenacil (Ref: DPX B634)*, <https://aeru.herts.ac.uk/aeru/ppdb/en/Reports/417.htm> (
